# Decoy diversification underpins the regulation of an NLR-mediated autoimmunity

**DOI:** 10.64898/2026.05.19.726385

**Authors:** Ye Jin Ahn, Naio Koehler, Jaepil Lee, Haseong Kim, Junhyeong Kim, Hwanjong Kim, Wanhui Kim, Min-Sung Kim, Young Jin Kim, Lennart Wirthmueller, Johannes Stuttmann, Alex Schultink, Kee Hoon Sohn

## Abstract

Appropriate activation of innate immune receptors is vital for the plant immune system. Immune activation must be strong and robust upon invasion by rapidly evolving pathogen species, while staying inactive in the absence of a pathogen to avoid constitutive defense responses that inhibit plant growth. Nucleotide-binding leucine-rich repeat receptors (NLRs) are intracellular proteins that surveil for pathogen invasion, often by direct or indirect perception of pathogen effector proteins. Indirect recognition of pathogen effector proteins, through monitoring the integrity of guardee or decoy proteins, can allow for a single NLR to detect a wide range of pathogen effectors. In this work, we investigated how the Solanaceae NLR NbPtr1 can recognize six different effector proteins from the bacterial pathogens *Pseudomonas syringae*, *Xanthomonas perforans*, and *Ralstonia pseudosolanacearum*. We identified several NOI-domain containing proteins that are guarded by NbPtr1. These NOI proteins share homology with, but are distinct from, RPM1 INTERACTING PROTEIN 4 (RIN4), a well-known guardee in Arabidopsis. Virus-induced gene silencing and CRISPR/Cas9-assisted deletion of *NbNOI* genes resulted in stunted growth dependent on NbPtr1 activity. Negative regulation of NbPtr1 requires the interaction between NbPtr1 and the suppressor NbNOIs. A conserved threonine residue of the NOI proteins is required for this interaction and fits into a putative binding pocket of NbPtr1 based on protein modeling. This threonine residue is modified by some of the recognized effector proteins. Our study uncovers the regulatory mechanism of an autoactive NLR and highlights the importance of decoy diversification in establishing compatibility with NLRs.

## INTRODUCTION

The plant immune system is specifically activated in response to pathogens through a sophisticated, multilayered pathogen detection system. Pathogens are recognized by pattern recognition receptors (PRRs) at the cell surface and nucleotide-binding leucine-rich repeat receptors (NLRs) inside the cell^1,2^. When PRRs bind to immunogenic microbial ligands, known as pathogen-associated molecular patterns (PAMPs), pattern-triggered immunity (PTI) is activated^1^. In turn, pathogens deliver specialized virulence proteins, called effectors, into host cells to suppress host immunity and promote virulence. In some cases, NLRs can recognize the specific effectors and activate effector-triggered immunity (ETI)^1^. Notably, ETI is often accompanied by a calcium influx and hypersensitive response, a type of programmed cell death (PCD) that prevents the spread of biotrophic pathogens at the site of infection^1,2^. PTI and ETI pathways are interconnected and mutually potentiate one another, thereby establishing robust host immunity^2–4^.

Coevolution between hosts and pathogens has resulted in diverse mechanisms of effector recognition by NLRs. Some NLRs indirectly detect pathogens by monitoring the host proteins targeted by effectors^5,6^. These host targets are called “guardees” if they have an alternative function independent of effector recognition, or “decoys” if they are mimics of effector virulence targets and used primarily for pathogen recognition^5,7,8^.

RPM1 INTERACTING PROTEIN 4 (AtRIN4) from *Arabidopsis thaliana* (hereafter, Arabidopsis) is a well-studied guardee that mediates multiple bacterial effector recognition by two Arabidopsis NLRs, RESISTANCE TO P. SYRINGAE 2 (RPS2) and RESISTANCE TO P. SYRINGAE PV MACULICOLA 1 (RPM1)^9–11^. AtRIN4 functions as a true guardee; it negatively regulates PTI, thus serving as a key target for pathogen effectors to suppress host immunity^12–15^. A defining structural feature of AtRIN4 is the presence of two nitrate-induced (NOI) domains at the N- and C-termini^15^. Although NOI domains were identified for nitrate-induced genes, a functional link between NOI-domain containing proteins and nitrogen metabolism remains elusive^16^. The NOI domains are about 40 amino acids long and harbor diverse post-translational modification sites, and effector-mediated modification of these sites triggers ETI^15^. For instance, a *Pseudomonas syringae* pv. *tomato* cysteine protease AvrRpt2 cleaves AtRIN4 at two sites in each NOI domain and interferes with AtRIN4 negative regulation of RPS2 autoactivity^10,17,18^. Meanwhile, AvrRpm1 from *P. syringae* pv. *maculicola* and AvrB from *P. syringae* pv. *glycinea* induce phosphorylation on AtRIN4 by recruiting RIPK (RPM1-induced protein kinase), and phosphorylation on Thr166 residue of AtRIN4 activates RPM1^19,20^. Later studies revealed that ADP-ribosylation by AvrRpm1 and rhamnosylation by AvrB on AtRIN4 could contribute to RPM1 activation^21,22^. Moreover, YopJ family effectors, which include HopZ5 from *P. syringae* pv. *actinidiae* and AvrBsT from *Xanthomonas euvesicatoria*, acetylate the same Thr166 residue and activate RPM1-dependent immunity^23^.

Significantly, AtRIN4-modifying effectors can target additional proteins that carry the NOI domain in Arabidopsis. AvrRpt2 can cleave NOI proteins at the AvrRpt2 cleavage site^17,18^, and AvrRpm1 induces ADP-ribosylation on ten NOI proteins other than AtRIN4^21^. These observations led us to hypothesize that NOI proteins function as effector targets that trigger ETI. Given that RIN4 is critical to PTI signaling^12,13^, NLRs likely evolved to guard a broader NOI protein family. The conservation of RIN4 and NOI proteins across land plants suggests that investigating the NOI protein family will uncover unique effector recognition strategies by NLRs in diverse taxa^16,24,25^.

To unravel the role of the NOI protein family in ETI, we investigated the diversified NOI proteins in *Nicotiana benthamiana*. In Solanaceae, AtRIN4-targeting effectors are recognized by a coiled-coil NLR called Pseudomonas tomato race 1 (Ptr1)^26–28^. It was shown that *N. benthamiana* homolog of Ptr1 (NbPtr1) has an expanded effector recognition specificity, as it can recognize AvrRpt2, AvrRpm1, AvrB, HopZ5 from *Pseudomonas*, AvrBsT from *Xanthomonas*, and RipBN, RipE1 from *Ralstonia*^27–29^. Overexpression of NbPtr1 in *N. benthamiana* causes cell death, suggesting that NbPtr1 is autoactive and might be negatively regulated in the absence of pathogens^27,28^. Previously, the involvement of RIN4 proteins in the suppression of Ptr1 autoactivity was hypothesized and tested with transient activity assays in *Nicotiana glutinosa* using *RIN4* orthologs from tomato and the *Ptr1* gene from *Solanum lycopersicoides*^27^. While suppression activity was observed, other experiments suggested that RIN4 orthologs may not be primary inhibitors of Ptr1. Notably, silencing the three *N. benthamiana RIN4* orthologs (*NbRIN4*) did not cause autoimmunity or have an observed effect on NbPtr1 function^28^. Based on this, we hypothesized that a more distantly related NOI protein, rather than a direct RIN4 ortholog, serves as the primary guardee/decoy of NbPtr1.

In this study, we identified several *N. benthamiana* NOI (NbNOI) proteins that are distinct from NbRIN4 orthologs and were able to suppress NbPtr1 autoactivity. Silencing the suppressor *NbNOI* genes in *N. benthamiana* caused lethality that was dependent on NbPtr1 activity. NbPtr1 suppression required direct interaction with the suppressor NbNOIs, and this interaction was disrupted in the presence of the effectors. The NbPtr1 autoactivity suppression mechanism was dissected at the amino acid residue level using AlphaFold3-guided structural investigation, followed by protein-protein interaction studies. Given that Ptr1 is not an ortholog of RPM1 or RPS2, our study demonstrates that NLRs have independently evolved to guard NOI proteins, which serve as convergent effector targets. We propose that NOI protein diversification generates novel decoys, providing a convergent mechanism for NLRs to detect diverse pathogen effectors.

## RESULT

### Several sequence-related NbNOIs suppress NbPtr1 autoactivity

To identify the primary guardee/decoy of NbPtr1, we performed a genome-wide search for NOI proteins in *N. benthamiana*. Three NbRIN4 orthologs and 20 more distantly related NbNOIs were identified from the search (Fig. 1A, Table S1, See Methods for details). Characteristic RIN4 features, such as AvrRpt2 cleavage site ([VIL]P[KRP]FGxWD motif), 2^nd^ RIN4 specificity motif (RSM2 to be Phe or Tyr), Thr166 corresponding residue (Thr or Ser), and a palmitoylation site at the C-terminus, are used as criteria for selecting NOI proteins^19,20,30,31^. Interestingly, the domain structures and amino acid sequence lengths of the NbNOIs were variable (Fig. 1A, Table S1).

**Figure 1.**
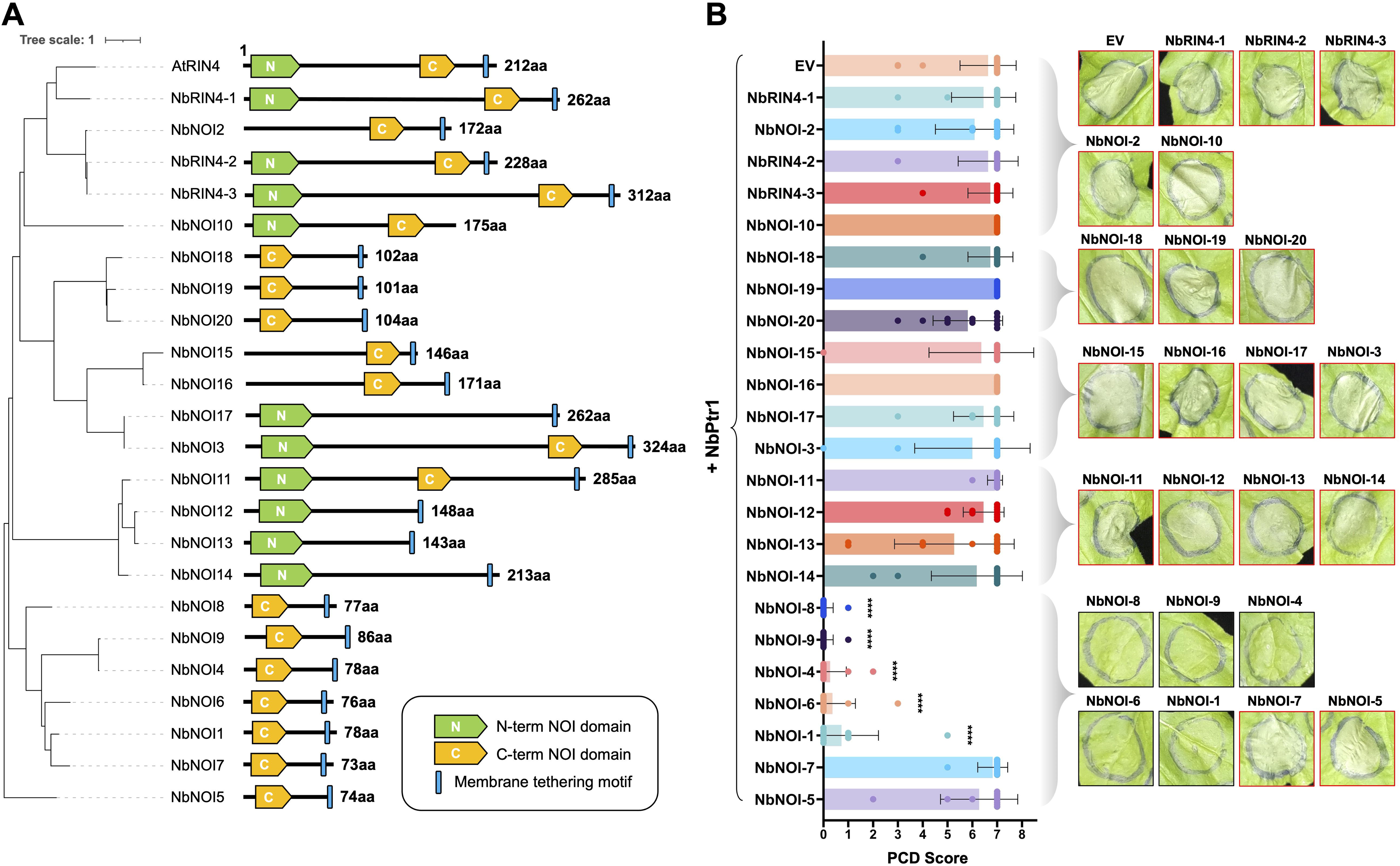
Several sequence-related NbNOIs suppress NbPtr1 autoactivity. (A) The phylogenetic tree of Arabidopsis RIN4 and NOI domain-containing proteins in *Nicotiana benthamiana*. The NOI proteins identified from 341 proteomes were used for reconstructing the phylogenetic tree (See Methods for details). To focus on the species of interest, non-*N. benthamiana* species sequences were removed while maintaining the original topology. The domain architecture of each protein is displayed with the phylogenetic tree. The protein size is indicated for each NOI protein. NOI domains are annotated as either N-NOI (Green) or C-NOI (Yellow) based on their sequence alignment with AtRIN4, and blue bars highlight the presence of membrane-tethering motifs. (B) NbPtr1 is suppressed by NbNOI-1, NbNOI-4, NbNOI-6, NbNOI-8, and NbNOI-9. On the first day of infiltration, EV, NbRIN4 orthologs, and NbNOIs (OD_600_ = 0.4) were expressed with P19 suppressor of post-transcriptional gene silencing (PTGS) (OD_600_ = 0.1) in Nb-1 *ptr1* using agroinfiltration. At 24 hpi, NbPtr1 (OD_600_ = 0.2) was expressed at the same infiltration sites. PCD was photographed at 4 dpi, and the photograph is a representative of three independent repeats with three to four technical repeats. A border surrounding each photograph was colored according to the mean PCD score: black (PCD < 2) or red (PCD > 5). On the left side of the photographs, a bar graph with the mean PCD score + SD is shown. For the statistical test, GRAPHPAD PRISM v.10.1.0 was used to perform the Kruskal-Wallis test followed by Dunn’s multiple comparisons test (*P* < 0.05). The comparison was made between the mean ranks of PCD scores for NbPtr1 + NbRIN4s/NbNOIs and PCD scores for NbPtr1 + EV. PCD graph bars without asterisks suggest no significant difference, and asterisks indicate significant differences: ****, *P* < 0.0001.

Due to its autoactive nature, we hypothesized that NbPtr1 would be negatively regulated in the absence of the corresponding avirulence effector recognition. Thus, we tested 23 *N. benthamiana* NOI family proteins, including three NbRIN4 orthologs, for the suppression of NbPtr1-induced PCD in the Nb-1 *ptr1* knockout line using *Agrobacterium*-mediated transient expression (hereafter, agroinfiltration) (Fig. 1B). Due to strong PCD induced by NbPtr1, EV and NbNOIs were expressed one day prior to NbPtr1 expression to ensure sufficient expression of NbNOIs (Fig. 1B, Table S2). NbNOI-1, NbNOI-4, NbNOI-6, NbNOI-8, and NbNOI-9 strongly suppressed NbPtr1-induced PCD, while the other NbNOIs and NbRIN4 orthologs did not reduce PCD development (Fig. 1B, Table S2). The five suppressor NbNOIs are phylogenetically closely related small proteins that carry only one C-NOI domain (Fig. 1A). All NbNOIs from the same clade were well expressed at the protein level (Fig. S1), confirming that the lack of cell death was not due to instability of NbNOI proteins.

### Silencing suppressor *NbNOIs* causes NbPtr1-dependent stunted growth phenotype

To further investigate NbPtr1 negative regulation by the suppressor NbNOIs, we silenced the suppressor *NbNOIs* using virus-induced gene silencing (VIGS) in Nb-1 wild-type and Nb-1 *ptr1*^28^ (Fig. 2A, 2B). In Nb-1 wild-type, silencing the suppressor *NbNOIs* resulted in stunted growth, compared to the growth observed in EV-VIGS control plants. Because *NbNOI-1* and *NbNOI-6* or *NbNOI-4* and *NbNOI-9* are nearly identical variants, VIGS was designed to silence similar genes together. The growth penalty was much stronger in *NbNOI-8*-VIGS plants than in *NbNOI-1,6*-VIGS or *NbNOI-4,9*-VIGS plants (Fig. 2A, 2B). Silencing *NbNOI-8* and additional *NbNOIs* caused lethality, as observed in *NbNOI-1,6,8*-VIGS, *NbNOI-4,8,9*-VIGS, and *NbNOI-1,4,6,8,9*-VIGS (Fig. 2A, 2B). The stunted growth phenotype observed in Nb-1 wild-type was not observed in Nb-1 *ptr1* (Fig. 2A, 2B).

**Figure 2.**
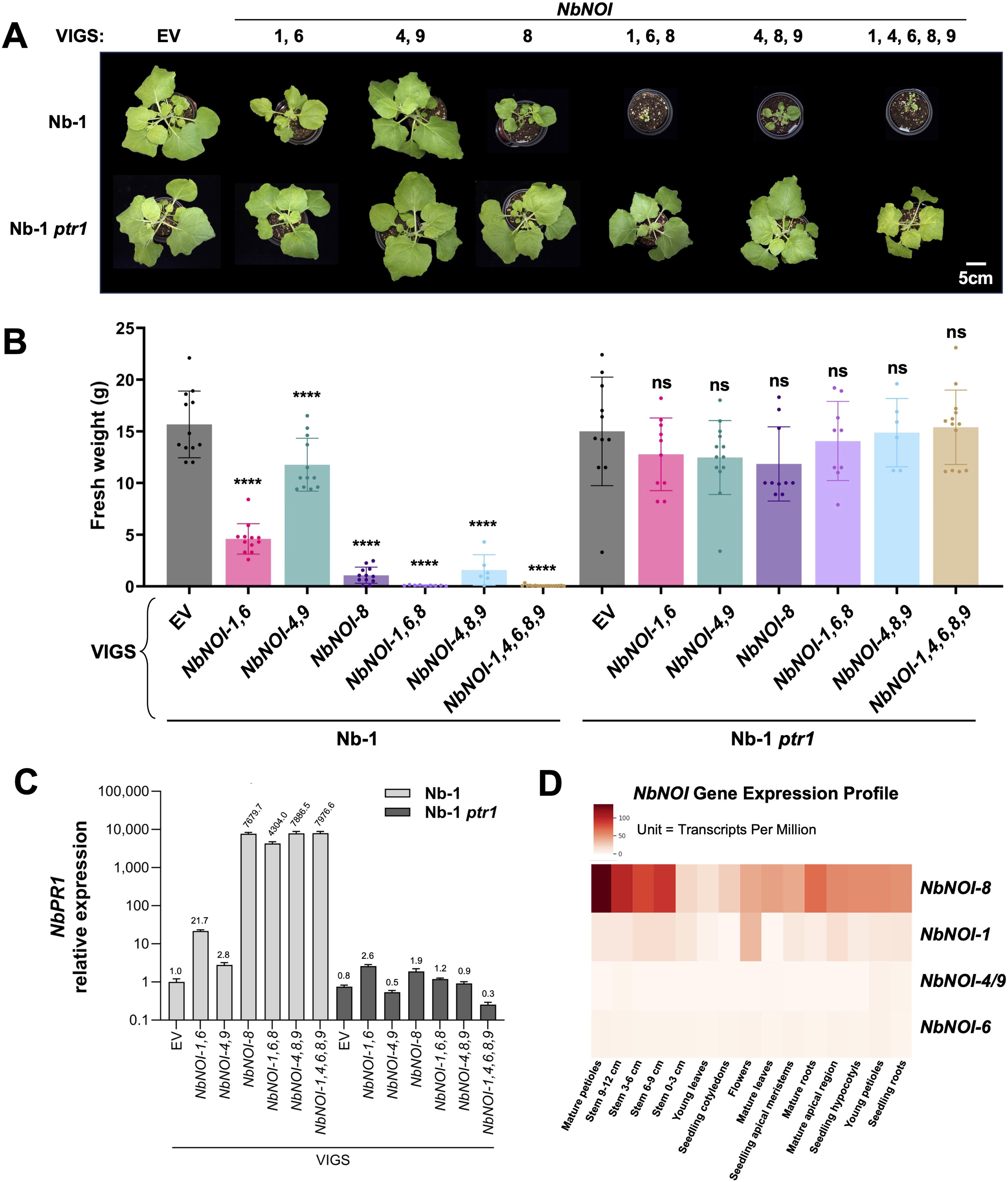
Silencing *NbNOIs* results in elevated defense, dependent on *NbPtr1* activity. (A) The stunted growth phenotype of *NbNOIs*-silenced plants is NbPtr1-dependent. Nb-1 and Nb-1 *ptr1* were silenced by VIGS. EV-VIGS plants were silenced with *lacZ* gene-carrying pTRV2 as a negative control. 8-week-old EV or *NbNOIs*-silenced plants were photographed from a bird’s-eye view. (B) Fresh weight measurement of EV or *NbNOIs*-silenced plants. 8-week-old silenced Nb-1 or Nb-1 *ptr1* were sampled above the hypocotyl, and the mass was measured. The fresh weight measurement was performed independently three times with three to six technical repeats. An ordinary one-way ANOVA test followed by Dunnett’s multiple comparison test was performed to compare each silenced plant’s weight to the EV-VIGS silenced plant’s weight for either Nb-1 or Nb-1 *ptr1* background. “ns” indicates no significance compared to EV-VIGS plants, while the asterisks signify significant differences: ****, *P* < 0.0001. (C) *NbPR1* gene expression is upregulated in *NbNOIs*-silenced Nb-1 plants. Two to four biological replicates of 7-week-old EV or *NbNOIs*-silenced Nb-1 and Nb-1 *ptr1* plants were sampled to measure *Nicotiana benthamiana PR1* gene expression by qRT-PCR. *NbPR1* expression of *NbNOIs*-silenced plants relative to EV-VIGS Nb-1 plants is shown in the graph. *NbPR1* expression was measured in three independent batches of silenced plants, and the representative data are shown in the figure. (D) NbNOI-8 has the highest expression level among the suppressor NbNOIs. The gene expression profiles of *NbNOIs* were searched and visualized as a heatmap using NbenBase browser^32^. The gene identifiers for *NbNOIs* according to NbenBase genome browser are as follows: NbNOI-1 (Nbe.v1.1chr10g14460), NbNOI-4 or NbNOI-9 (Nbe.v1.1.chr16g07390), NbNOI-6 (Nbe.v1.1chr15g45400), and NbNOI-8 (Nbe.v.1.1chr13g15100).

Total RNA was extracted from silenced plants to assess silencing efficiency and defense marker gene expression (Fig. S2, 2C). The expressions of *NbNOI-1* and *NbNOI-8* were reduced in the silenced plants, suggesting that the stunted growth was caused by the reduced expression of the *NbNOIs* (Fig. S2). The expression of *NbNOI-4* or *NbNOI-9* was challenging to determine, since a high Ct value (> 34) was measured consistently in Nb-1 and Nb-1 *ptr1* EV-VIGS plants during RT-qPCR, suggesting that the native expression levels of *NbNOI-4* and *NbNOI-9* are comparably low (Fig. S2). Furthermore, *NbPR1* expression was increased only in the *NbNOI*s silenced plants in Nb-1 wild-type, but not in Nb-1 *ptr1* (Fig. 2C). Notably, any Nb-1 plants silenced with *NbNOI-8* showed 4,000∼8,000 times higher expression of *NbPR1* than the expression measured in EV-VIGS (Fig. 2C). According to *N. benthamiana* gene expression profile^32^, *NbNOI-8* is expressed most strongly among the suppressor *NbNOIs* (Fig. 2D), explaining why the stunted growth was strongest with the *NbNOI-8*-VIGS construct.

Using CRISPR-Cas9-mediated gene editing, *nbnoi1,6* double and *nbnoi1,6 nbptr1* triple knock-out lines were generated in the Nb-0 background to test the negative regulation of NbPtr1 by NbNOI-1 and NbNOI-6 (Fig. S3). A minor growth defect was observed in *nbnoi1,6*, and *nbnoi1,6 nbptr1* showed improved growth (Fig. S4), consistent with the NbPtr1-dependent stunted phenotypes of *NbNOIs* silenced plants (Fig. 2). Additionally, NbPtr1-recognizing effectors were agroinfiltrated to Nb-0 wild-type, *nbnoi1,6* double, and *nbnoi1,6 nbptr1* triple knock-out lines (Fig. S5, Table S3). All of the effectors triggered strong PCD in *nbnoi1,6* double knock-out line, similarly to the response observed in Nb-0 wild-type (Fig. S5, Table S3). In *nbnoi1,6 nbptr1* triple knock-out line, AvrRpt2, RipE1, AvrRpm1, AvrB-triggered PCD was abolished, but HopZ5 and AvrBsT-triggered PCD was only partially reduced due to the additional recognition by NbZAR1^28^ (Fig. S5, Table S3). Despite knocking out *NbNOI-1* and *NbNOI-6*, strong effector-triggered cell deaths were observed in *nbnoi1,6* double mutant. In summary, the five suppressor NbNOIs redundantly function for the negative regulation of and effector recognition by NbPtr1. Our several attempts to generate *nbnoi1,4,6,8,9* pentuple mutant were unsuccessful, likely due to the lethal phenotype observed in *NbNOI-1,4,6,8,9*-VIGS plants (data not shown).

### Effectors disrupt NbNOIs-mediated suppression of NbPtr1

Effector-triggered cell death observed in *nbnoi1,6* double knock-out line strongly suggested that the suppressor NbNOIs may redundantly function for NbPtr1-mediated immunity (Fig. S5). However, generating five *NbNOIs* knockout plants or using five *NbNOIs* silenced plants is challenging, since knock-out or silenced plants were lethal (Fig. 2). To test if the suppressor NbNOIs redundantly function for effector recognition, we agroinfiltrated effectors under conditions in which NbPtr1 is suppressed by NbNOIs and monitored changes in the suppression phenotype (Fig. 3A, 3B). We VIGSed *NbNOI-1,4,6,8,9* in Nb-1 *ptr1* plants to avoid endogenous *NbPtr1* and *NbNOIs* gene expression during the experiment. The synthesized NbNOI-1, NbNOI-4, and NbNOI-8 (SynNOI-1, SynNOI-4, and SynNOI-8) with altered codons were used to prevent gene silencing when transiently expressed in *NbNOI-1,4,6,8,9*-VIGS plants (Table S4). EV and SynNOIs were expressed one day before the expression of NbPtr1 with GFP or the corresponding effectors (Table S5). Co-expression of NbPtr1, GFP, and SynNOIs did not induce PCD since NbPtr1 was suppressed by SynNOIs (Fig. 3A, 3B, Table S6). Importantly, co-expression of AvrRpt2, RipE1, AvrRpm1, AvrB, HopZ5, or AvrBsT instead of GFP induced robust PCD, indicating that these effectors interfere with NbNOI-mediated suppression of NbPtr1 autoactivity. AvrRpm1 induced weak PCD, consistent with previous observations^28^. In summary, we showed that all tested effectors interfere with NbNOIs-mediated suppression of NbPtr1 and induce cell death.

**Figure 3.**
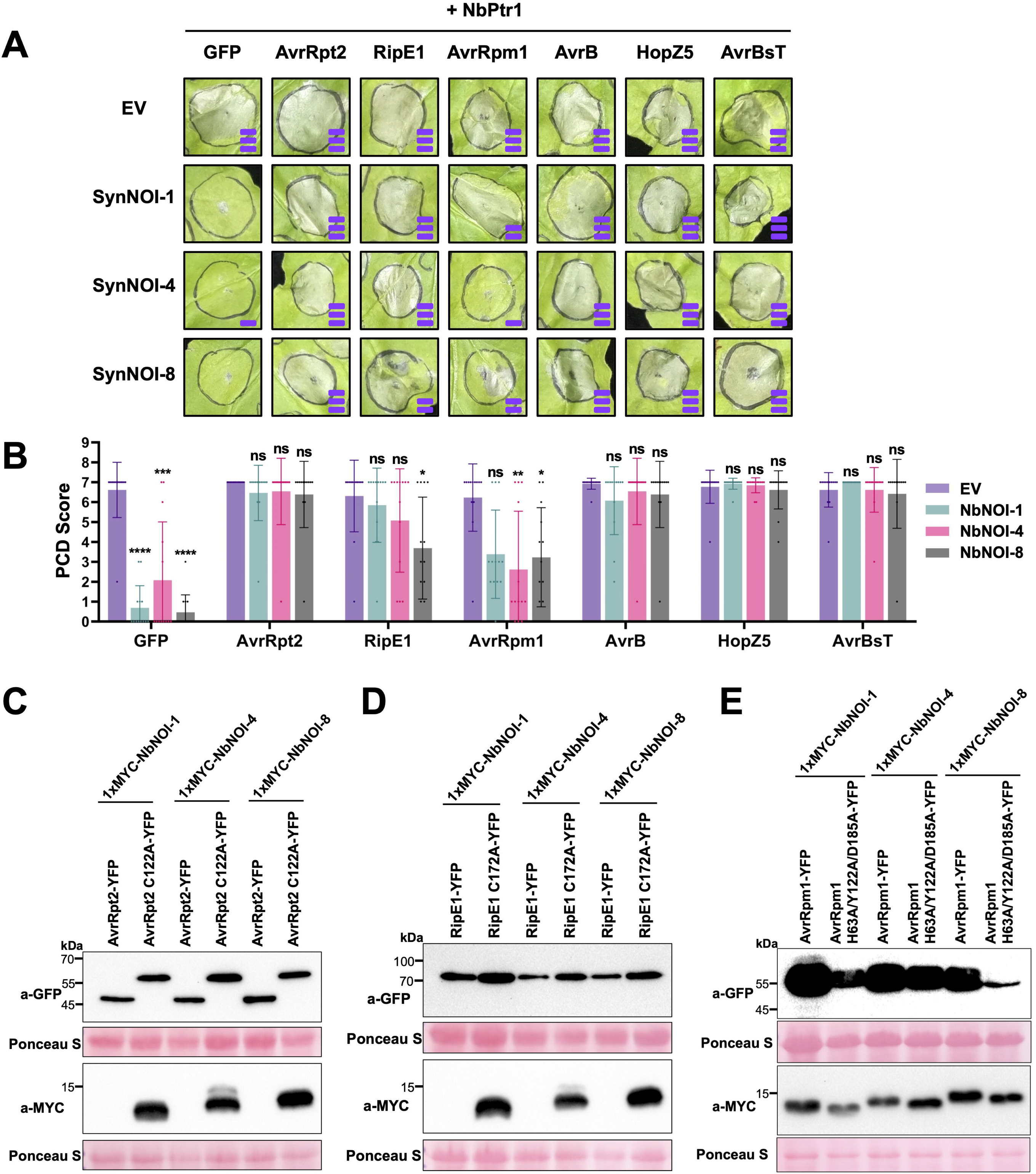
Effectors interfere with NbNOI-directed suppression of NbPtr1-autoactivity. (A) The additional expression of effectors disrupts NbPtr1 suppression by NbNOIs. In Nb-1 *ptr1 NbNOI-1,4,6,8,9*-VIGS plants, EV and codon-optimized SynNOIs were agroinfiltrated at OD_600_ = 0.4 with P19 (OD_600_ = 0.1). The next day, NbPtr1 (OD_600_ = 0.2) with GFP, AvrRpt2, RipE1, AvrRpm1, AvrB, HopZ5, AvrBsT (OD_600_ = 0.4) were coinfiltrated at the same infiltration spots. The PCD photographs are from 4 to 7 dpi and are representative of three independent repeats, each with four to five technical repeats. PCD intensity is indicated by purple bars stacked at the bottom right corner of each photograph. The mean PCD score is categorized as follows: 0-1 (no bar), 1-3 (one bar), 3-5 (two bars), and 5-7 (three bars). (B) PCD scoring of Fig. 3A. A bar graph of the mean PCD score + SD is shown. GRAPHPAD PRISM v.10.1.0 was used to perform the Kruskal-Wallis test followed by Dunn’s multiple comparisons test (*P* < 0.05) to test if the mean rank of PCD scores of NbPtr1 + SynNOI + Effector (or GFP) is significantly different from the mean rank of PCD scores of NbPtr1 + EV + Effector (or GFP). “ns” indicates no significance from the statistical test. Asterisks indicate significant differences: *, *P* < 0.1; **, *P* < 0.01; ***, *P* < 0.001; ****, *P* < 0.0001. (C) AvrRpt2 degrades NbNOI-1, NbNOI-4, and NbNOI-8. AvrRpt2-YFP and AvrRpt2 C122A-YFP were coexpressed with 1xMYC tagged NbNOI-1, NbNOI-4, and NbNOI-8 (all OD_600_ = 0.4) with P19 (OD_600_ = 0.1) using agroinfiltration in Nb-1 *ptr1*. (D) RipE1 degrades NbNOI-1, NbNOI-4, and NbNOI-8. RipE1-YFP and RipE1 C172A-YFP were coinfiltrated with 1xMYC tagged NbNOI-1, NbNOI-4, and NbNOI-8 (all OD_600_ = 0.4) with P19 (OD_600_ = 0.1) using agroinfiltration in Nb-1 *ptr1*. (E) AvrRpm1 causes a band shift to NbNOI-1, NbNOI-4, and NbNOI-8. AvrRpm1-YFP and AvrRpm1 H63A/Y122A/D185A-YFP were coinfiltrated with 1xMYC tagged NbNOI-1, NbNOI-4, and NbNOI-8 (all OD_600_ = 0.4) with P19 (OD_600_ = 0.1) using agroinfiltration in Nb-1 *ptr1*. For the anti-MYC blot, 47 μg of protein samples were loaded to observe clear band shifts caused by AvrRpm1.

### Effectors induce post-translational modifications of NbNOIs

The effectors recognized by NbPtr1 induce diverse post-translational modifications on AtRIN4^10,15,17–19,21–23^. Notably, the five suppressor NbNOIs harbor most of the residues that correspond to the key residues in AtRIN4 that were shown to be directly or indirectly modified by the effectors (Fig. S6). We hypothesized that NbNOIs would undergo similar modifications as observed in AtRIN4 due to the significant level of sequence conservation.

Among the reported diverse post-translational modifications of AtRIN4, proteolytic cleavage by cysteine proteases AvrRpt2 and RipE1 can be detected by Western blot. Due to the high sequence identity between NbNOI-1 and NbNOI-6 (as well as NbNOI-4 and NbNOI-9), we selected NbNOI-1, NbNOI-4, and NbNOI-8 for Western blotting. N-terminally 1xMYC-tagged NbNOI-1, NbNOI-4, and NbNOI-8 were co-expressed with C-terminally YFP-tagged AvrRpt2 or its catalytic mutant AvrRpt2 C122A^33^ in Nb-1 *ptr1* (Fig. 3C). The NbNOIs were not detected in the anti-MYC blot from the samples co-expressed with AvrRpt2 wild-type, but were strongly detected from the samples co-expressed with AvrRpt2 C122A (Fig. 3C). As shown previously, the AvrRpt2 cleavage site is located in close proximity to the N-terminus of NbNOIs (near amino acid 15) (Fig. S6), making it difficult to detect the processed NbNOI N-terminal fragment in immunoblot analysis due to its small size^17,18^.

Similarly, NbNOI-1, NbNOI-4, and NbNOI-8 were not detected when co-expressed with RipE1, unlike their strong expressions when co-expressed with RipE1 C172A mutant^29,34^ (Fig. 3D). RipE1 is a putative cysteine protease that does not share significant sequence identity with AvrRpt2 and does not cleave itself when expressed in plant cells^29,34^ (Fig. 3D, S7A). N-terminally 1xMYC-tagged NbNOI-1 F11A is resistant to AvrRpt2-directed cleavage^17^ (Fig. S7B). Interestingly, RipE1 could not cleave the NbNOI-1 F11A mutant, indicating that RipE1 and AvrRpt2 may have overlapping target specificity (Fig. S7B).

Soybean RIN4s and AtRIN4 co-expressed with AvrRpm1 were previously shown to exhibit reduced mobility when detected by immunoblot^21^. NbNOI-1, NbNOI-4, and NbNOI-8 were co-expressed with AvrRpm1 or AvrRpm1 H63A/Y122A/D185A mutant^21,35^ in Nb-1 *ptr1* (Fig. 3E). NbNOIs co-expressed with AvrRpm1 showed band shifts, while co-expression with AvrRpm1 catalytic mutant did not (Fig. 3E). The band shifts on NbNOIs are consistent with the ADP-ribosylation of NbNOIs by AvrRpm1. In summary, these results demonstrate that the effectors can target and induce modifications of the NbNOIs that suppress NbPtr1 autoactivity. The modifications are similar to those induced by these effectors on AtRIN4 in plant cells.

### NbPtr1 interacts specifically with the suppressor NbNOIs

To test if NbPtr1 and NbNOIs physically associate with each other, we set out a co-immunoprecipitation (co-IP) assay using transiently expressed NbPtr1 and NbNOIs (Fig. 4A). EV and 1xMYC-NbNOIs were co-expressed with NbPtr1-6xHA in Nb-1 *ptr1* (Fig. 4A). NbPtr1 co-IPed with NbNOI-1, NbNOI-4, and NbNOI-8 but not with NbNOI-5, demonstrating that the interaction was specific between NbPtr1 and the suppressor NbNOIs. Even though NbNOI-5 is closely related to the suppressor NbNOIs compared to other NbNOIs (Fig. 1A), the lack of interaction between NbPtr1 and NbNOI-5 may have caused the inability to suppress NbPtr1 (Fig. 1, 4A).

**Figure 4.**
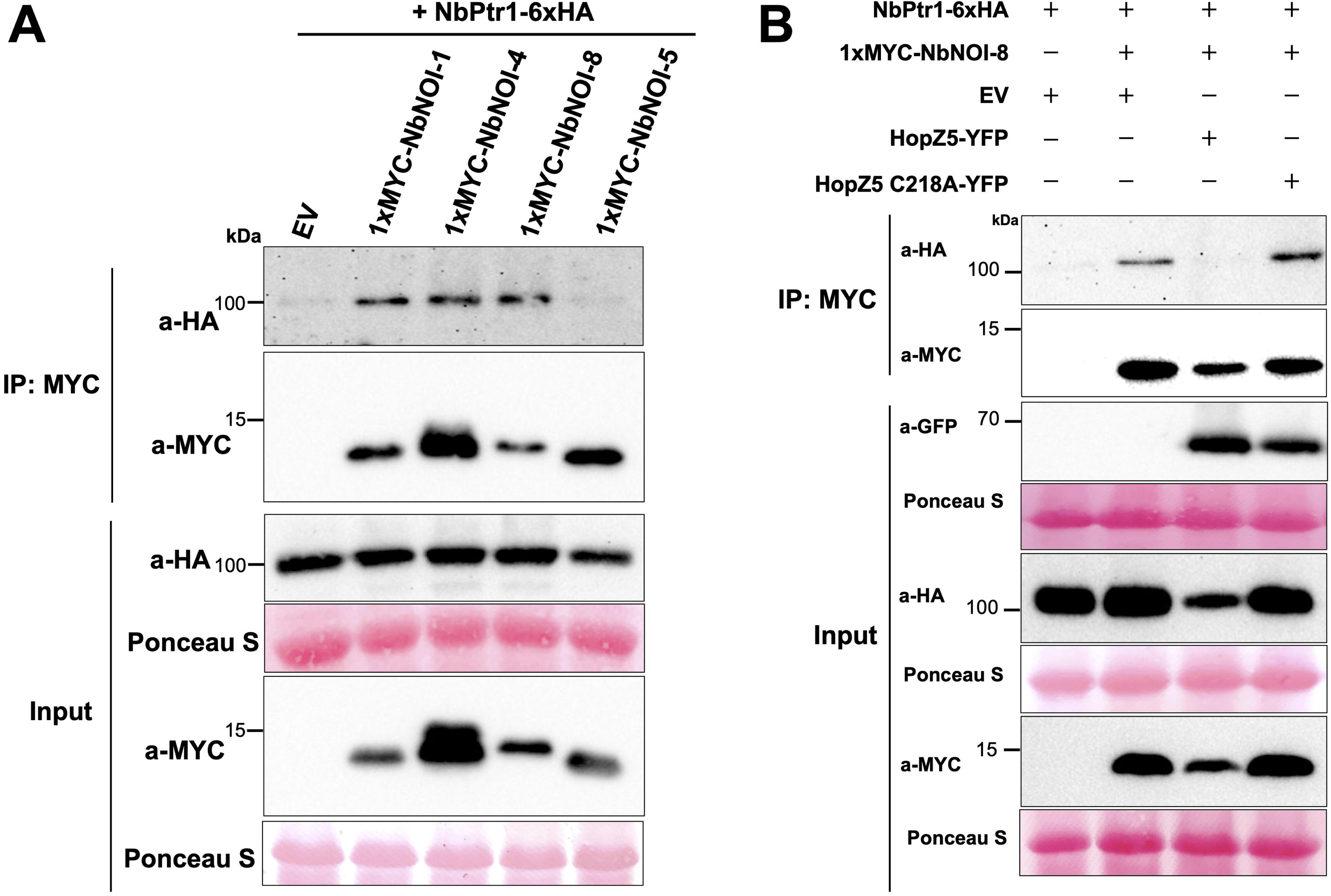
NbPtr1 specifically interacts with the suppressor NbNOIs. (A) NbPtr1 is co-IPed with NbNOI-1, NbNOI-4, NbNOI-8 but not with NbNOI-5. NbPtr1-6xHA (OD_600_ = 0.05) was expressed with EV and 1xMYC tagged NbNOI-1, NbNOI-4, NbNOI-8, and NbNOI-5 (OD_600_ = 0.4) using agroinfiltration in Nb-1 *ptr1*. Co-IP was performed independently three times, and the representative result is shown in the figure. _(B)_ HopZ5 weakens the interaction between NbPtr1 and NbNOI-8. NbPtr1-6xHA (OD_600_ = 0.05), 1xMYC-NbNOI-8, EV, HopZ5-YFP, and HopZ5 C218A (OD_600_ = 0.4) were coexpressed using agroinfiltration in NbZAR1-VIGS Nb-1 *ptr1* plants. Co-IP was performed independently three times, and the representative result is shown in the figure.

We further investigated whether the NbPtr1-NbNOI interaction is altered in the presence of an effector. EV, NbPtr1-6xHA, 1xMYC-NbNOI-8, C-terminally YFP tagged HopZ5 and HopZ5 C218A^36^ were agroinfiltrated in Nb-1 *ptr1* knock-out plants silenced for *NbZAR1* (Fig. 4B). Nb-1 *ptr1 NbZAR1*-VIGS plants were used, since HopZ5 is recognized by both NbPtr1 and NbZAR1^28^. Significantly, NbPtr1 was not co-IPed with NbNOI-8 in the presence of wild-type HopZ5, whereas HopZ5 C218A mutant variant did not alter the NbPtr1-NbNOI8 association. This indicates that effector-mediated disruption of NbPtr1-NbNOI association activates immunity. We did not investigate if disruption of NbPtr1 association with NbNOIs recruits helper NLRs, since *NLR required for cell death* (*NRC*) genes were not required for NbPtr1 function (Fig. S8, Table S7)^34,37^. Moreover, mutations of hydrophobic residues (IVV to EEE) in the N-terminal a1 helix eliminated NbPtr1 autoactivity, suggesting that NbPtr1 may activate through oligomerization, similarly to singleton NLRs such as ZAR1^38,39^ (Fig. S9, Table S8). Taken together, NbPtr1 and NbNOI association is disrupted upon effector-directed modification of NbNOIs that may induce NbPtr1 oligomerization and activate NRC-independent cell death.

### The conserved threonine residue of NbNOIs is crucial for the negative regulation of NbPtr1

To better understand the structural basis for NbNOI-NbPtr1 association, we used AlphaFold3 to predict the heterodimeric structure of NbPtr1 and the NOI domain (8-37 aa) of NbNOI-8 (Fig. 5A). The predicted structure showed that the NOI domain of NbNOI-8 fits within the NB-ARC (nucleotide-binding domain shared by Apaf-1, R proteins, and CED4)^40^ and LRR domains of NbPtr1. Interestingly, a cavity-like structure was observed inside the ARC2 domain of NbPtr1 (Fig. S10A). NbNOI-8 had a protruding structure formed by the conserved threonine (Thr26 for NbNOI-8), which corresponds to Thr166 of AtRIN4, and the threonine fits right inside the cavity of NbPtr1, analogous to a ball-and-socket interaction (Fig. S10B). The cavity structure was formed by Glu431, Pro433, and Val477 (“EPV”) residues of NbPtr1, and these residues were consistently predicted to interact with the conserved threonine residue of the five suppressor NbNOIs that corresponds to Thr166 of AtRIN4 (Fig. 5B, S10, Table S9). In this model, Glu431 forms a hydrogen bond with the hydroxyl group of Thr26 of NbNOI-8, and Pro433 and Val477 provide hydrophobicity that increases the interaction via the methyl group of Thr26 of NbNOI-8 (Fig. 5B).

**Figure 5.**
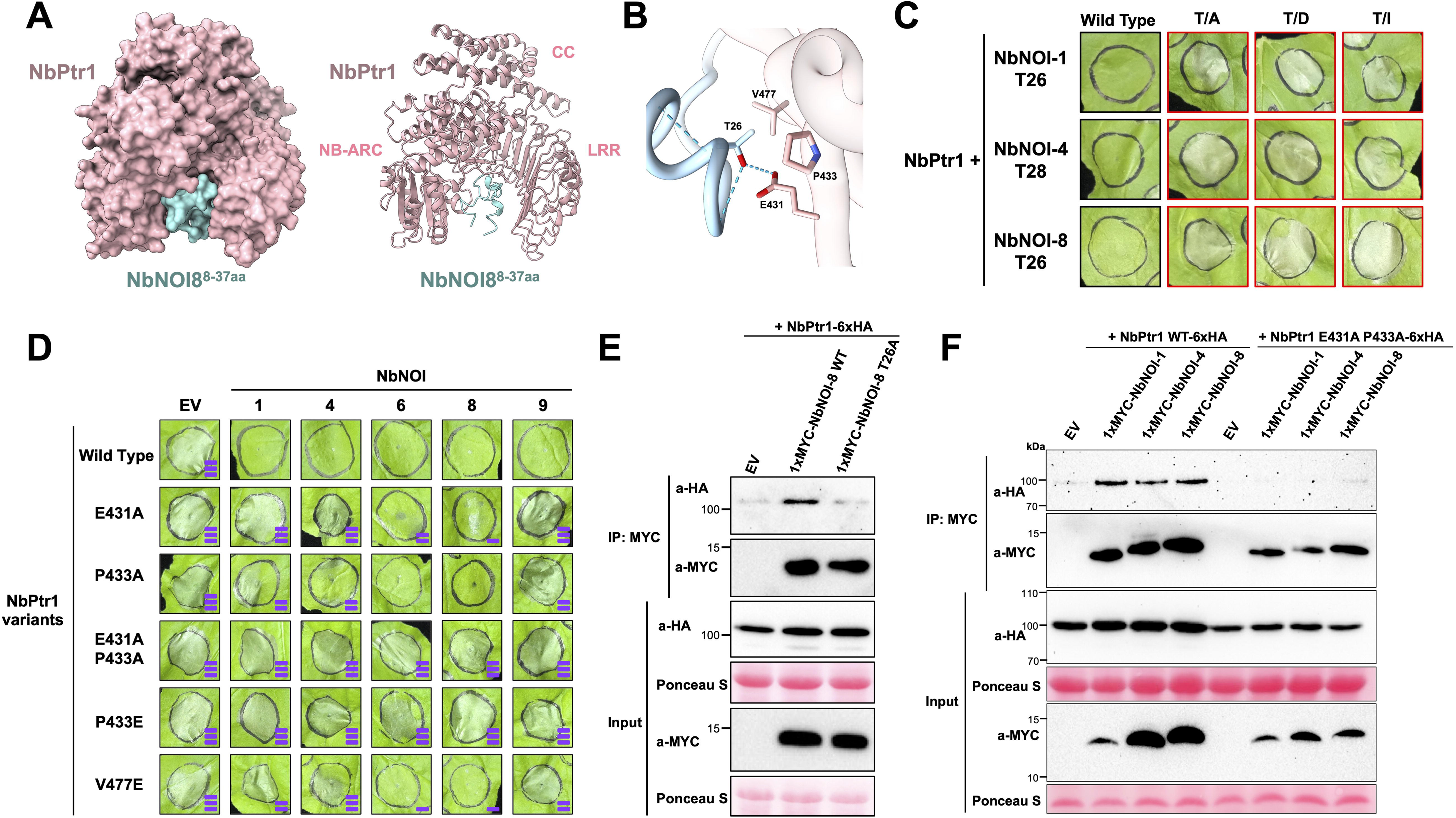
The conserved threonine residue of NbNOIs is critical for the negative regulation of and interaction with NbPtr1. (A) AlphaFold3 prediction of the interacting structure of NbPtr1 and NbNOI-8 NOI domain (8-37 aa). The interacting structure of NbPtr1 and NbNOI-8 NOI domain was predicted using a seed value of 2 (pTM = 0.81; ipTM = 0.43). (B) Three key residues of NbPtr1 interact with Thr26 of NbNOI-8. The close-up view of NbPtr1 interaction with the conserved threonine residue of NbNOI-8 is demonstrated in the figure. NbPtr1 residues that are within 6Å distance of Threonine 26 of NbNOI-8 are highlighted. Glu431 forms a hydrogen bond with Threonine 26 of NbNOI-8, which is indicated with a blue dotted line. (C) The conserved threonine residue of NbNOIs is critical for NbPtr1 suppression. On the first day of infiltration, NbNOI-1, NbNOI-4, NbNOI-8 wild-type and mutants at OD_600_ = 0.4 with P19 (OD_600_ = 0.1) in Nb-1 *ptr1*. The next day, NbPtr1 (OD_600_ = 0.2) was expressed in the same infiltration spots, and PCD was observed and photographed at 4 dpi. Representative photographs of three independent repeats, each with four biological replicates, are shown in the figure. A border surrounding each photograph was colored according to the mean PCD score: black (PCD < 2) or red (PCD > 5). (D) EPV residues of NbPtr1 are critical for negative regulation by NbNOIs. EV and NbNOIs were expressed at OD_600_ = 0.4 with P19 (OD_600_ = 0.1) in Nb-1 *ptr1*. The next day, NbPtr1 variants were expressed at OD_600_ = 0.2 on the same infiltration sites. PCD was photographed and scored at 4 dpi, and the photographs shown in the figure represent results from three independent repeats with three to four biological replicates. PCD intensity is indicated by purple bars stacked at the bottom right corner of each photograph. The mean PCD score is categorized as follows: 0-1 (no bar), 1-3 (one bar), 3-5 (two bars), and 5-7 (three bars). (E) Mutation of NbNOI-8 Thr26 to Ala disturbs the interaction with NbPtr1. NbPtr1-6xHA (OD_600_ = 0.05) was coexpressed with EV, 1xMYC-NbNOI-8 wild-type, and 1xMYC-NbNOI-8 T26A (OD_600_ = 0.4) in Nb-1 *ptr1*. Co-IP was performed independently three times, and the representative result is shown in the figure. (F) NbPtr1 E431A P433A does not interact with NbNOI-1, NbNOI-4, and NbNOI-8. NbPtr1 wild-type-6xHA and NbPtr1 E431A P433A-6xHA (OD_600_ = 0.05) were coexpressed with EV, 1xMYC-NbNOI-1, 1xMYC-NbNOI-4, and 1xMYC-NbNOI-8 (OD_600_ = 0.4) in Nb-1 *ptr1*. Co-IP was performed independently three times, and the representative result is shown in the figure.

To test the significance of the conserved threonine residue of NbNOIs for NbPtr1 suppression, the threonine residues of NbNOI-1,4,8 were mutagenized to alanine (null mutant; T/A), aspartate (phosphomimic mutant; T/D)^19^, and isoleucine (acetylmimic mutant; T/I)^23^ and tested for changes in their suppression ability (Fig. 5C, S11A, Table S10). Mutations at the threonine residue (T/A, T/D, and T/I) of any tested NbNOIs could not suppress NbPtr1 autoactivity (Fig. 5C, S11A, Table S10). NbNOIs and their threonine mutant variants were well-expressed at the protein level, indicating that the conserved threonine residue is indispensable for the suppression of NbPtr1 autoactivity (Fig. S11B).

The residues in the putative binding pocket of NbPtr1 were mutated to test their significance in NbNOI-mediated suppression of NbPtr1 autoactivity (Fig. 5D). Glu431, Pro433, and Val477 were mutated to alanine, and Pro433 and Val477 were additionally mutated to glutamate to become negatively charged. When expressed alone, all NbPtr1 mutant variants maintained the autoactivity comparable to NbPtr1 wild-type (Fig. S12, Table S11). The five suppressor NbNOIs failed to suppress most of the cell deaths induced by NbPtr1 mutant variants (Fig. 5D, S13, Table S12). NbNOI-6 and NbNOI-8 could weakly suppress PCD induced by NbPtr1 P433A and V477E. However, NbPtr1 E431A P433A and NbPtr1 P433E induced PCD consistently, regardless of which suppressor NbNOI was coexpressed, as shown from the PCD scoring and the conductivity measurement (Fig. 5D, S13, Table S12). The suppressor NbNOIs and NbPtr1 wild-type and mutant variants were well-expressed at the protein level (Fig. S14), suggesting that the lack of cell death suppression was not due to the instability of NbNOI protein variants.

Based on our cell death assay results (Fig. 5C, 5D), we hypothesized that the constitutive activation of NbPtr1 is due to the lack of physical association with suppressor NbNOIs. Therefore, we tested whether the alanine mutation on the conserved threonine residue of NbNOI-8 (T26A) affects the interaction with NbPtr1 (Fig. 5E). NbNOI-8 T26A showed significantly reduced interaction with NbPtr1 as compared to wild-type NbNOI-8, indicating that the Thr26 residue is critical for interaction with NbPtr1. Additionally, the NbPtr1 E431A P433A mutant variant, which NbNOIs could not suppress, was subjected to co-IP with NbNOIs. In contrast to wild-type NbPtr1, NbPtr1 E431A P433A mutant variant was not co-IPed with any of the NbNOIs (Fig. 5F). These results strongly demonstrate that effector-induced post-translational modifications of NbNOIs destabilize their interactions with autoactive NLR NbPtr1.

## DISCUSSION

In this study, we investigated the diversified family of NOI proteins and their role in immune signaling in plants. We identified several NbNOIs with a single NOI domain that suppress NbPtr1 autoactivity. We also show that multiple bacterial pathogen effectors target and modify NbNOIs post-translationally, thereby interfering with the suppression of NbPtr1 autoactivity. AlphaFold3-guided structural study elucidated that the interaction between the conserved threonine residue of NbNOIs and “EPV” residues of NbPtr1 is critical for the suppression and activation of immunity. We propose a coevolutionary model in which NOI protein duplication yielded novel decoys that suppress NbPtr1 autoactivity and facilitate effector sensing, allowing NbPtr1 to specialize toward these decoys rather than the ancestral virulence targets (Fig. 6).

**Figure 6.**
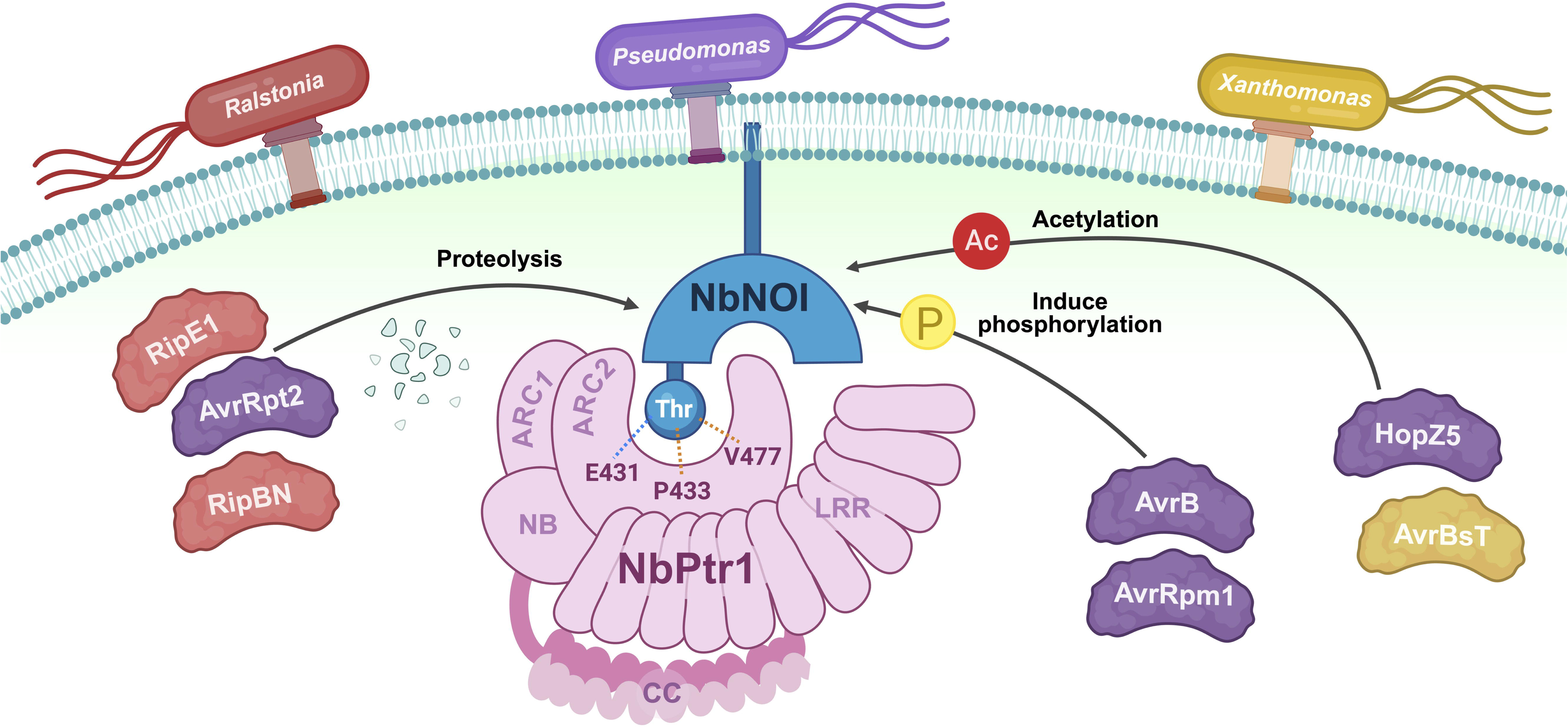
NbPtr1 negative regulation by suppressor NbNOIs. Among the diversified family of NOI domain-containing proteins, NbNOI-1, NbNOI-4, NbNOI-6, NbNOI-8, and NbNOI-9 negatively regulate NbPtr1 through direct interaction. The conserved threonine residue of NbNOIs that corresponds to AtRIN4 Thr166 interacts primarily with Glu431, Pro433, and Val477 residues from ARC2 of the NB-ARC domain of NbPtr1. Their interaction is analogous to a ball-and-socket interaction, with the conserved threonine residue sitting right inside the cavity formed by the three residues from NbPtr1. The interaction between NbNOIs and NbPtr1 is key to the negative regulation of NLR-mediated autoimmunity. NbNOI-NbPtr1 interaction is disrupted by bacterial effectors with diverse biochemical activities. Post-translational modifications by the effectors disrupt the interaction between NbNOIs and NbPtr1, thereby inducing ETI.

Here, our study characterizes previously unknown members of the NOI family as suppressors of an autoactive NLR. NbNOIs with a single NOI domain play major roles in suppressing NbPtr1 and inducing ETI. The NOI family display diverse protein structures with either one or two NOI domains (Fig. 1A), and this variation in the number of NOI domains is observed from mosses to flowering plants^24^. Despite the sequence and structural diversity, two NOI domains-carrying RIN4 has been most extensively studied in this family due to its functional importance as a guardee^15^. Interestingly, in Arabidopsis, AtRIN4 fragments produced by AvrRpt2, known as AvrRpt2 Cleavage Products (ACPs), have been shown to participate in PTI suppression and RPS2-mediated immunity^12,41,42^. The active AtRIN4 fragments include the soluble ACP2 carrying the N-NOI domain and the membrane-tethered ACP3 carrying the C-NOI domain of AtRIN4. Analogous to the suppressor NbNOIs, ACP3 suppresses RPS2, whereas ACP2 overrides the suppression and activates RPS2^42^. Consequently, the studies on AtRIN4 active fragments and the suppressor NbNOIs demonstrate that a single NOI domain is sufficient to achieve NLR regulation. This observation raises two fundamental questions: What drives the diversification of the NOI protein family and how is the specific compatibility of an NOI protein and a guard NLR established?

We hypothesize that the diversified forms of the NOI protein family likely arose from coevolution with guard NLRs. Given that RIN4 and NOI proteins have important roles in PTI^12,13^, pathogen effectors frequently target these proteins to suppress host defense^16,17,21,24^. To counteract this virulence strategy, plants evolved NLRs to monitor NOI proteins. Following *NOI* gene duplication, the resulting paralogs may have undergone functional divergence: one copy maintains its ancestral role in PTI^5^, while the other evolves into a specialized decoy. This duplication allows the NLRs to coevolve exclusively with the decoy, eventually no longer recognizing the original virulence target. Because these decoys are liberated from the functional constraints as PTI suppressors, they can rapidly evolve to optimize effector recogntion^8^. Ultimately, the diverse forms of NOI proteins emerged from enhancing compatibility with their respective NLRs, while maintaining structural characteristics essential for interaction with effectors. The observation that non-orthologous NLRs, such as NbPtr1, RPM1, RPS2, and FB_MR5, monitor diverse forms of NOI proteins suggests that NOI proteins serve as convergent guardees/decoys for NLRs^25^.

NOIs may have primarily diversified to coevolve with guard NLRs, but their diversification provides functional redundancy that aids in effector recognition. All five suppressor NbNOIs can effectively suppress NbPtr1-induced PCD and mediate effector recognition (Fig. 1B, 3A, S5). Moreover, NOI diversity causes differences in gene expression level and tissue-specific expression pattern (Fig. 2D). For example, NbNOI-8 expresses highly in most tissues, and NbNOI-1 expresses specifically in flowers (Fig. 2D). These differences among suppressor NbNOIs may help to complement each other to regulate NbPtr1 in different tissues or environmental conditions. It is also plausible that unknown effectors target specific suppressor NbNOIs; in which scenario, having multiple NbNOIs as decoys would benefit plants. Notably, most NLRs require a specific decoy to recognize effectors, or if they associate with multiple decoys, each decoy is specialized to recognize different effectors, as observed in the ZAR1-RLCK (receptor-like cytoplasmic kinase) network^25,43^. In contrast, NbPtr1 immune activation demonstrates a unique mechanism in which multiple decoys share the identical function in effector recognition.

Several studies have shown that *N. benthamiana* RIN4 orthologs are involved in NbPtr1-mediated immunity. RipE1 has been shown to interact and colocalize with NbRIN4 at the plasma membrane^44,45^. NbRIN4-1 and kiwifruit RIN4 (AcRIN4-2) were shown to interact with NbPtr1, and NbPtr1 and AcRIN4-2 interaction was disrupted by HopZ5^46^, similar to our finding where the suppressor NbNOI-NbPtr1 interaction is disrupted by HopZ5 (Fig. 4B). It was also reported that weak cell death induced by the coiled-coil (CC) domain of NbPtr1 was suppressed partially by NbRIN4, and the CC domain of NbPtr1 interacts with NbRIN4^44^. However, silencing NbRIN4 orthologs did not cause any growth defect or altered effector-triggered cell death^28,44^. In contrast, we observed severe lethality and elevated defense responses when the suppressor NbNOIs were silenced in *N. benthamiana* (Fig. 2). Based on our data presented in this study, we propose that the five suppressor NbNOIs identified in our study, rather than NbRIN4 orthologs, are key players in the activation of NbPtr1-mediated immunity.

How do plants negatively regulate autoactive NLRs? Negative regulators of plant immunity are crucial, as inappropriate activation of immunity can severely compromise plant survival^47,48^. Oftentimes, autoimmunity is caused by the absence or modification of a guardee that is surveilled by an NLR^47,48^. For instance, *saul1-1* mutant displays SOC3-dependent autoimmunity in Arabidopsis, where SAUL1 (Senescence-Associated E3 Ubiquitin Ligase I) is guarded by SOC3 (Suppressor of *chs1-2, 3*; TNL) and its TN partners^49,50^. Similarly, knocking out MAP kinase cascade genes (*mekk1*, *mkk1 mkk2*, *mpk4*) causes autoimmunity activated by SUMM2 (Suppressor of *mkk1 mkk2* 2; CNL)^51–54^. HopAI1 from *Pseudomonas* inactivates the MAP kinase cascade, and perturbations on MAPKs activate SUMM2-mediated immunity, accompanied by resistosome ring-like structures consisting of EDS1-PAD4-ADR1-L1^52,55,56^. As these examples show, a guardee and a guard NLR often physically interact, and this interaction may be key to regulating autoimmunity^57–59^. Notably, most studies on autoimmunity focus on the mechanistic relationship between NLRs and immune signaling components but offer little information on cognate effectors and the signals that activate the NLRs. Our study on suppressor NbNOIs and NbPtr1 demonstrates autoimmunity regulation through direct interaction, as well as how this regulation is essential for effector recognition. The conserved threonine residue of NbNOIs is required for NbPtr1 suppression (Fig. 5) and also serves as a hotspot for modifications by effectors (Fig. 3, S6). Since the effectors and NbPtr1 share the same threonine residue for their mode of action, effector recognition occurs in tandem with NLR activation. Our findings suggest that the interaction interface between a guardee and its NLR defines the physical site where effectors target the guardee. Consequently, this site can be leveraged to find novel candidate effectors recognized by NLRs. As demonstrated here, a structure-guided approach not only elucidates the molecular basis of the guardee-NLR association but also provides the framework to determine whether autoimmunity and ETI converge on the same downstream signaling pathways.

Modification of the conserved threonine residue on NbNOIs is a key for activating NbPtr1, but NbPtr1 recognizes effectors that do not necessarily modify the threonine of NbNOIs. NbPtr1 recognizes cysteine proteases, including AvrRpt2, RipBN, and RipE1^27–29^, which cause cleavage after glycine in the PxFGxW motif of the NOI domain, and this does not directly modify the conserved threonine residue (Fig. S7). In addition, AvrRpm1 ADP-ribosylates Asp185 of GmRIN4b^21^, and the corresponding residue in NbNOIs is glutamate, a few residues apart from the threonine (Fig. S6). It is possible that proteolysis and ADP-ribosylation change the conformation of NbNOIs, preventing the conserved threonine from interacting with EPV residues of NbPtr1, or there could be additional residues other than the threonine that participate in NbPtr1 suppression. Likewise, most NbNOIs, including NbNOI-5, which is phylogenetically closest to the suppressor NbNOIs, have the conserved threonine residue but cannot suppress or interact with NbPtr1. The NbNOI-5 NOI domain and NbPtr1 interaction model is not significantly different to that of the suppressors, which suggests that sequence polymorphisms outside the NOI domain may stabilize the interaction. Therefore, the conserved threonine residue is necessary but insufficient to confer compatibility with NbPtr1. Further investigation of the additional residues will provide a comprehensive understanding of NbNOI-NbPtr1 compatibility.

NbPtr1’s ability to recognize multiple sequence-unrelated effectors originated from several bacterial pathogens is desirable for engineering disease-resistant crops. In commercial tomato, *Ptr1* is a pseudogene that lacks a start codon, so there has been an effort to introgress functional *Ptr1* from a wild relative of tomato, *Solanum lycopersicoides,* into a commercial tomato variety^26,27^. Remarkably, *S. lycopersicoides* introgression lines were maintained heterozygous for the *Ptr1* locus, because homozygous *Ptr1/Ptr1* individuals were rarely found and grew more slowly than other genotypes^27^. Based on our study, this phenomenon may be caused by a lack or insufficient expression of functional NOI proteins that can negatively regulate Ptr1. This highlights the importance of transferring decoys that can properly regulate NLR to resolve restricted taxonomic functionality. Recently, co-transfer of a Solanaceae sensor and helper NLRs into distantly related plant species has successfully conferred disease resistance^60^. Taken together, the co-transfer of Ptr1 and suppressor NOI would be a desirable strategy for the successful development of bacterial disease-resistant crops.

In conclusion, we unveil the function of the diversified NbNOIs as suppressors of the autoactive NLR, NbPtr1. NOI proteins likely diversified and formed compatibility with diverse NLRs, since they play a substantial role in basal immunity and effector recognition. We anticipate that further exploration will reveal unknown functions of NOI proteins in ETI, supporting our proposal that the NOI protein family represents a convergent guardee/decoy system that coevolves with guard NLRs. Our findings position the NOI family as a central hub for decoy diversification and immune signaling, offering a new perspective on how NLRs sense diverse pathogen effectors.

## METHODS

### Plant materials and growth conditions

*Nicotiana benthamiana* Domin was grown at 22 C in a short-day condition (11 h light and 13 h dark per day). Hanarum soil mix (64.3% cocopeat, 15% peatmoss, 2.5% vermiculite, 10% perlite, 8% zeolite; Goesan, Chungcheongbuk-do, South Korea) was used to grow *N. benthamiana*. Nb-1 and Nb-0 backgrounds of *N. benthamiana* were used, and the difference between the two lines is determined by the expression level of JIM2 (XOPJ4 IMMUNITY 2)^61^. JIM2 is an RLCK required for the recognition of XopJ4, HopZ5, and AvrBsT by NbZAR1^28,61,62^. Since Nb-1 has a lower expression level of JIM2 than Nb-0, Nb-1 was used to study NbPtr1 without NbZAR1 recognition^28^.

### Bacterial strains growth and transformation

*Escherichia coli* DH5α and *Agrobacterium tumefaciens* AGL1 and GV3101 strains were grown in low-salt Luria Broth (LB) liquid or agar (20 g/L) media with appropriate antibiotics (Duchefa Biochemie, Haarlem, Netherlands). For bacterial transformation, electrocompetent *E. coli* and *A. tumefaciens* cells were pulsed with plasmids (1.8 kV for *E. coli* and 2.0 kV for *A. tumefaciens*). The transformed cells were mixed with LB liquid medium and incubated at 37C for an hour for *E. coli* and 28C for two hours for *A. tumefaciens.* Transformed cells were plated on LB agar plates and incubated at their optimal temperature until colonies were observed.

### NOI protein search and phylogenetic tree reconstruction

In search of NOI proteins in *N. benthamiana*, protein sequences from two different *N. benthamiana* genomes^63,64^ were extracted. The initial search was performed to find proteins with the [V, I, L]P[K, R, P]FGxWD motif. Among the candidate proteins, those lacking the 2^nd^ RIN4 specificity motif [F, Y] and T166 residue [T, S] were eliminated. Furthermore, proteins that lack a palmitoylation motif [C, S, F] (two to four repeats) at the C-terminal were eliminated. As a result, three NbRIN4 orthologs^65^ and 20 NbNOI proteins were identified (Table S1). Following this approach, 4,936 NOI proteins were retrieved from proteomes of 341 plant species, mostly consisted of angiosperms^66^. Sequences were aligned using MAFFT (v7.525)^67^ and subsequently trimmed using trimAl (v1.5)^68^ to remove gap-rich regions. A maximum-likelihood phylogenetic tree was reconstructed using FastTreeMP (v2.2.0)^69^. To focus on the *N. benthamiana* family within an evolutionary context, non-*N. benthamiana* sequences were pruned, while the original tree topology using the ETE3 toolkit (v3.1.3)^70^. The final phylogeny of AtRIN4 and NbNOI proteins was visualized using iTOL (v7; https://itol.embl.de/)^71^.

### Plasmid construction

Sequences of NbNOIs, SynNOIs, NbNOI-4 T28A/D/I, NbNOI-8 T26 A/D/I, and NbPtr1 EEE were synthesized (Twist BioScience, South San Francisco, CA, USA) and ligated into pUC19b for Golden Gate (GG) assembly^72^. NbNOIs and NbRIN4-related sequences were assembled into pICH86966 with 35s promoter, N-terminal 1xMYC tag, and NOS terminator. NbPtr1 and its mutants were assembled into pICH86988 either with a C-terminal 6xHA tag or a 4xMYC tag. AvrRpt2, RipE1, AvrRpm1, AvrB, HopZ5, and AvrBsT were assembled to pICH86988 either with a C-terminal 6xHA tag or a YFP tag^28,29^. Information about the catalytic site mutants of the effectors, including AvrRpt2 C122A, RipE1 C172A, and AvrRpm1 H63A/Y122A/D185A, is described in previously published studies^28,29,33–35^. For VIGS constructs, NbNOI VIGS target sequences from Nb-0 cDNA were cloned into pUC19b and assembled into pTRV2-GG using GG assembly (Table S14). The NbZAR1-VIGS used in Fig. 4 is the same construct described in the previous publication^28^. For CRISPR vectors, oligonucleotides for sgRNAs were synthesized (Bioneer, Daejeon, South Korea) and hybridized into shuttle vectors (Table S14, S15)^73^. The shuttle vectors carrying sgRNAs were assembled into the CRISPR-Cas9 vector, pDGE687, using GG assembly, as previously described^73^ (Table S15). Information on the final CRISPR-Cas9 vectors is shown in Table S15.

### Site-directed mutagenesis

For site-directed mutagenesis of NbNOI-1 T26A/D/I, PCR was performed using PrimeSTAR GXL DNA Polymerase (Takara Bio Inc., Japan) with pUC19b carrying NbNOI-1 and primers that amplify the mutagenesis region (Table S14). The PCR product digested with DpnI, and the digested product was transformed into *E. coli* DH5α. In case of NbPtr1 cloning, NbPtr1 sequence was divided into four modules and cloned into pUC19b for GG assembly. For NbPtr1 E431A, P433A, E431A P433A, and P433E mutagenesis, we amplified the partial sequence of NbPtr1 with forward primers with mutations (Table S14), since E431 and P433 located at very 5’ end of NbPtr1 third module sequence. The PCR product was ligated into pUC19b using SmaI. For mutagenesis of NbPtr1 V477E and P-loop (K202R), NbPtr1 pUC19b module vectors (module 3 for V477E and module 2 for K202R) were amplified with mutagenesis primers (Table S14). The mutagenesis primers were flanked by BsaI recognition sites and four nucleotide overhangs to facilitate ligation during the subsequent process. PCR products were purified using the GeneAll PCR SV kit (GeneAll Biotechnology, South Korea) and digested with BsaI and DpnI. The digested products were purified with GeneAll PCR SV and ligated using T4 DNA ligase. The ligated products were transformed into *E. coli* DH5α. The pUC19b vectors carrying mutagenized sequences were used for GG assembly.

### Virus-induced gene silencing

*A. tumefaciens* carrying pTRV2-GG with the target gene sequence and pTRV1 were grown on LB agar media at 28C for one day. *A. tumefaciens* cells were cultured in LB liquid media and grown at 28C overnight. The saturated cultures were centrifuged to pellet the cells, and the cells were resuspended in the VIGS buffer (10 mM MgCl2, 1 mM MES (2-(N-morpholino)ethanesulfonic acid) [pH 5.6]). The resuspended cells were diluted to OD_600_ = 0.5, and *A. tumefaciens* carrying pTRV1 and pTRV2-GG were mixed at a 1:1 ratio. One-week-old *N. benthamiana* seedlings were inoculated with the *A. tumefaciens* VIGS mixture and grown in a short-day condition until the plants became six to seven weeks old, as described previously^28,74^.

### Gene expression analysis

Total RNA was isolated from *N. benthamiana* leaf disc using Trizol (Ambion, USA). RNA was treated with DNase I (Sigma, Germany) and used for cDNA synthesis with Maxima First strand cDNA synthesis kit for RT-qPCR (Thermo Fisher Scientific, Waltham, MA, USA). The RT-qPCR was performed in a 96-well plate using CFX-connect real-time system (BioRad, Hercules, CA, USA) with GoTaq qPCR master mix (Promega, USA). All RT-qPCR experiments were carried out with three biological replicates (independently harvested samples) and three technical replicates each. Gene expression levels were normalized to the reference gene *NbEF1α*. The oligonucleotide sequences used for RT-qPCR are listed in Table S14.

### Generation of N*. benthamiana* mutants using CRISPR/Cas9

Guide RNAs for target genes were designed using CRISPR-P v2.0 with the following criteria: PAM site as NGG, U6 promoter, 20-nucleotide-long guide sequence, and target genome *N. benthamiana* (v0.4.4). Guide sequences containing BsaI, BbsI, and polyT sites were removed. Guide sequences with the highest On-score and the fewest possible off-target sites were selected. Since *N. benthamiana* is allotetraploid, guide RNAs were selected to target both the target gene and its homolog, assuming that the homolog has a similar function. The guide sequences with ATTG/GTTT overhangs are listed in Table S14. The Cas9-expressing vectors with guide sequences were transformed into *A. tumefaciens* AGL1, which was used for transformation in *N. benthamiana* by Tomato Biotech (Jinju, Gyeongsang-namdo, South Korea). Genomic DNA of the transformed plants was extracted using the CTAB method and used for PCR with genotyping primers and gel electrophoresis (Table S14). DNA with appropriate sizes was extracted from the gel and sent for sequencing (NICEM, Seoul, South Korea; Macrogen, Seoul, South Korea). Mutations were analyzed using Geneious v 2025.2.1 (https://www.geneious.com) and TIDE v5.0.5 (Tracking of Indels by Decomposition; https://apps.datacurators.nl/tide/).

### *Agrobacterium*-mediated transient expression in *N. benthamiana*

*A. tumefaciens* was grown on LB agar media at 28C for one day. *A. tumefaciens* cells were cultured in LB liquid media and grown at 28C overnight. The saturated cultures were centrifuged to pellet the cells, and the cells were resuspended in the Agro buffer (10 mM MgCl2, 10 mM MES [pH 5.6]). The resuspended cells were diluted according to the OD_600_ information indicated in the figure legends. Five to six-week-old *N. benthamiana* was infiltrated with a needleless syringe. PCD was scored using a scale from 0-7 and statistically analyzed using GRAPHPAD PRISM v.10.1.0 (GraphPad Software, San Diego, CA, USA www. graphpad.com), as previously described^28^.

### Ion leakage measurement

Three leaf discs (d = 1cm) were sampled from each agroinfiltration in *N. benthamiana*. Three replicates (n = 3) of leaf discs were resuspended in 2 ml of deionized water for 2 hours with gentle shaking. The conductivity was measured using a LAQUAtwin-EC-33 compact conductivity meter (Horiba, Kyoto, Japan).

### Protein extraction

For quick protein extraction, six leaf discs (d = 1cm) per sample were sampled from agroinfiltrated *N. benthamiana* at 48 hpi and finely ground in liquid nitrogen. After grinding, 200 ul of the extraction buffer (250 mM Tris-Cl [pH 6.8], 8% sodium dodecyl sulfate, 0.1% bromophenol blue, 0.1% glycerol, 17 mM DTT) was added, and the samples were thawed for 10 min on ice and vortexed. The samples were boiled at 95C for 5 min and stored at -20C. For co-IP, one full leaf (around 1 g) of agroinfiltrated *N. benthamiana* at 48 hpi was sampled for each sample. The leaf samples were finely ground in liquid nitrogen using a mortar and a pestle. The extraction buffer for NbPtr1 (10% glycerol, 50 mM Tris, 2 mM EDTA, 150 mM NaCl, 10 mM DTT, 1x Sigma Complete^TM^ mini protease inhibitor (100x is one tablet diluted in 500ul deionized water), 0.2% NP-40, 0.5% Triton-X, 1 tablet of PhosSTOP^TM^ 1 tablet, 0.25% DDM) was added to the samples and further ground on ice until the samples thawed and became finely ground. The extracted samples were filtered through Miracloth (Millipore, Germany) and snap-frozen in liquid nitrogen for storage or used immediately for co-IP.

### Co-immunoprecipitation

20 ul/sample of EZview^TM^ Red Anti-c-Myc Affinity Gel (Millipore, Germany) was washed with the bead wash buffer (10% glycerol, 50 mM Tris, 2 mM EDTA, 150 mM NaCl, 10 mM DTT, 1x Sigma Complete^TM^ mini protease inhibitor, 0.2% NP-40, 0.5% Triton-X) three times (6,000 rpm for 30 sec). The washed beads were added to 1 mg of the extracted samples and incubated at 4C for 2 hours. The incubated beads were washed three times and boiled at 95C for 5 min in 30 ul of 2x Laemmli buffer containing 5% 2-Mercaptoethanol.

### SDS-PAGE and Western blot

Protein samples were loaded onto SDS-PAGE gels and run at 80V for 30 min and 100V for at least 1.5 hr. The gels were transferred to PVDF membranes at a constant 25V condition using the Trans-blot Turbo Transfer System (BioRad, Hercules, CA, USA). Transferred membranes were blocked with 5% skim milk in TBS/T for 30 min. The membranes were incubated with appropriate primary antibodies overnight at 4C. The membranes were washed three times for 7 min in TBS/T, then incubated with secondary antibodies for 1 hour at room temperature. The membranes were washed with TBS/T and visualized using SuperSignal West Pico, Dura, and Femto chemiluminescent substrates (Thermo Fisher Scientific, Waltham, MA, USA). Antibody information is as follows: Myc-Tag (9B11) Mouse monoclonal antibody (2276, Cell Signaling), GFP Polyclonal antibody (A-6455, Invitrogen), anti-HA (3F10, Roche), Goat anti-rabbit IgG-peroxidase (A0545, Sigma), Rabbit anti-mouse IgG-peroxidase (A9044, Sigma), and Goat anti-rat IgG-peroxidase (A9037, Sigma).

### Using AlphaFold3 to find NbPtr1-NbNOI interaction residues

NbPtr1 full-length and the NOI domain of NbNOIs protein sequences were submitted to the AlphaFold3 Server, and the protein complexes were predicted with three different seed values (1, 2, or 3) (Table S8). Interaction residues were identified in the predicted structures by using the interface function in ChimeraX 1.10. NbPtr1 residues that are in close proximity (maximum 6Å) with the conserved threonine residue were identified using the distance function in ChimeraX. The residues identified more than five times were selected for mutagenesis and suppression assays.

## ACKNOWLEDGEMENT

We thank Chih-Hang Wu (Academia Sinica, Taiwan) for providing *nrc234* knockout plants. We appreciate Jongyoon Park, Hokyun Yoo, and Gwanyeong Nam (Seoul National University, South Korea) for assisting in genotyping CRISPR knock-out plants. This project was supported by the National Research Foundation of Korea (NRF) grants (RS-2024-00463021, RS-2023-NR077218, RS-2025-00512558).

## AUTHOR CONTRIBUTIONS

Conceptualization: Y.J.A., A.S., K.H.S. Designed experiments: Y.J.A., K.H.S.

Performed experiments and analyzed: Y.J.A., N.K., J.P.L., H.S.K., J.H.K., H.J.K., W.H.K., Y.J.K.

Generated materials and resources: Y.J.A., H.S.K., A.S., J.S. Experimental insights: M.S.K., A.S., L.W., K.H.S.

Wrote the manuscript: Y.J.A., A.S., L.W., K.H.S. Financial support: K.H.S., Y.J.A.

---

**Supplementary Figure 1. NbNOI protein expression in *Nicotiana benthamiana*.**

N-terminally 1xMYC tagged NbNOI-1, NbNOI-4, NbNOI-6, NbNOI-8, NbNOI-9, NbNOI-5, and NbNOI-7 were expressed (OD_600_ = 0.4) with P19 (OD_600_ = 0.1) in Nb-1 *ptr1* using agroinfiltration.

**Supplementary Figure 2. *NbNOI* VIGS efficiency in silenced *N. benthamiana*.**

Two to four biological replicates of 7-week-old EV or *NbNOIs*-silenced Nb-1 and Nb-1 *ptr1* plants from Fig. 2A were sampled for total RNA extraction. Quantitative RT-PCR was performed on three independent batches of silenced plants, and the representative data are shown in the figure.

**Supplementary Figure 3. *nbnoi1,6* and *nbnoi1,6 nbptr1* knock-out lines mutation map**

Two guide RNAs were designed for each gene, *NbNOI-1*, *NbNOI-6*, and *NbPtr1,* to generate CRISPR knock-out mutants. For each gene, genomic DNA is presented with exons (yellow arrows) and introns (black line) to show the locations of sgRNAs. Guide RNA sequences are shown as green letters, and PAM sites are indicated as red. Any mutations in mutant lines are indicated in red in the sequence.

**Supplementary Figure 4. *nbnoi1,6* knock-out line shows NbPtr1-dependent minor growth defect.**

8-week-old Nb-0 wild-type, *nbnoi1,6* double, and *nbnoi1,6 nbptr1* triple knock-out lines were photographed from the front view and a bird’s-eye view. The size scale is indicated in the bottom-left corner.

**Supplementary Figure 5. NbNOIs redundantly function for the recognition of effectors.**

GFP, AvrRpt2, RipE1, AvrRpm1, AvrB, HopZ5, and AvrBsT are agroinfiltrated at OD_600_ = 0.4 with P19 (OD_600_ = 0.1) in Nb-0 wild-type, *nbnoi1,6* double, and *nbnoi1,6 nbptr1* triple knock-out lines. Programmed cell death (PCD) was photographed at 4-5 days post-infiltration (dpi), and representative photographs from three independent repeats are shown in the figure.

A border surrounding each photograph was colored according to the mean PCD score: black (PCD < 2), dotted red (2 < PCD < 5), or red (PCD > 5). Below the photographs, a bar graph of the mean PCD score + SD is shown. GRAPHPAD PRISM v.10.1.0 was used to perform the Kruskal-Wallis test, followed by Dunn’s multiple comparisons test (*P* < 0.05), to test if the mean rank of PCD scores from Nb-0 wild-type is significantly different from the mean rank of PCD scores from two other knockout lines. If the mean ranks of PCD scores from Nb-0 wildtype and mutant lines are not significantly different, “ns” is indicated in the graph. PCD graph bars are labeled with asterisks to indicate significant differences: ****, *P* < 0.0001; **, *P* < 0.01.

**Supplementary Figure 6. Protein sequence alignment of AtRIN4 and NbNOIs at the C-NOI domain**

Amino acid sequences of the C-terminal NOI domains of AtRIN4, NbNOI-1, NbNOI-4, NbNOI-5, NbNOI-6, NbNOI-7, NbNOI-8, and NbNOI-9 are aligned. Amino acid residue numbers are indicated on the left and right of the sequence alignment. AtRIN4 residues that are known to be modified post-translationally by effectors and induce effector-triggered immunity are indicated above each residue. AvrRpt2 recognizes the VPxFGxW motif on AtRIN4 and cleaves its target after Gly152, activating RPS2-mediated immunity^10,17^. AvrRpm1 can ADP-ribosylate its targets, and soybean RIN4 was shown to be ADP-ribosylated at Asp153^21^. AvrRpm1 and AvrB induce phosphorylation on the conserved Thr166 of AtRIN4, activating RPM1^19,20^. A recent study suggested that AvrB functions as a rhamnosylator, adding UDP-rhamnose on Thr166 of AtRIN4^22^. HopZ5 and AvrBsT acetylate AtRIN4, and the acetylation on Thr166 activates RPM1^23^. RIN4 specificity motifs (RSM1 and RSM2) that are important for regulating RPS2, RPM1, and FB_MR5 are indicated as well^30^.

**Supplementary Figure 7. RipE1 cannot degrade AvrRpt2 cleavage-resistant NbNOI-1 F11A mutant.**

(A) Protein sequence alignment at the AvrRpt2 cleavage site. Amino acid sequences of the effectors used in this study with protease activities (RipE1, AvrRpt2, and RipBN), AtRIN4, and NbNOIs that can suppress NbPtr1 are aligned at the AvrRpt2 cleavage site (RCS). RipE1 is a putative cysteine protease, and RipBN is a *Ralstonia* homolog of AvrRpt2. The conserved VPxFGxW motif is highlighted in blue, and any unmatched sequence is indicated with darker shades of blue. Amino acid residue numbers are indicated on the left and right of the sequence alignment. AtRIN4 contains two NOI domains with RCS; therefore, RCS1 and RCS2 are both present in the alignment. The black arrow indicates a potential cleavage site based on previous studies of AvrRpt2 and AtRIN4^10,17,18^.

(B) AvrRpt2, RipBN, and RipE1 cannot degrade NbNOI-1 F11A mutant. The conserved Phe11 in the RCS region of NbNOI-1 was mutated to alanine to be resistant to AvrRpt2 cleavage activity. N-terminally 1xMYC tagged NbNOI-1 and NbNOI-1 F11A mutant (OD_600_ = 0.4) were coexpressed with C-terminally 6xHA tagged AvrRpt2, AvrRpt2 C122A, RipBN, and RipE1 (OD_600_ = 0.4) with P19 (OD_600_ = 0.1) in Nb-1 *ptr1*. The expected sizes of AvrRpt2-6xHA (28.4 kDa), AvrRpt2 C122A-6xHA (35.6 kDa), RipBN-6xHA (27 kDa), and RipE1-6xHA (53.3 kDa) are indicated as red asterisks on the anti-HA blot.

**Supplementary Figure 8. NbPtr1-recognized effectors induce NRC-independent cell death.**

PCD induced by the effectors recognized by NbPtr1 is NRC independent. EV, AvrPto + Pto, Rpi-blb2 + Avrblb2, AvrRpt2, RipE1, AvrRpm1, AvrB, HopZ5, and AvrBsT were expressed at OD_600_ = 0.4 in *Nicotiana benthamiana* Nb-0 wild-type and *nrc234*. P19 (OD_600_ = 0.1) was added to all infiltrations. PCD was scored and photographed at 3 dpi, and representative photographs from three independent repeats are shown in the figure. A border surrounding each photograph was colored according to the mean PCD score: black (PCD < 2) or red (PCD > 5). The mean PCD score + and SD are shown in the bar graph. GRAPHPAD PRISM v.10.1.0 was used to perform the Mann-Whitney test (*P* < 0.05) to test if the mean rank of PCD scores from Nb-0 wild-type is significantly different from the mean rank of PCD scores from *nrc234*. If the mean ranks of PCD scores from Nb-0 wildtype and *nrc234* are not significantly different, “ns” is indicated in the graph. PCD graph bars are labeled with asterisks to indicate significant differences: ****, *P* < 0.0001.

**Supplementary Figure 9. NbPtr1 P-loop motif and N-terminal hydrophobic residues are essential for autoactivity.**

(A) Protein sequence alignment at the MADA motif region of AtZAR1, NbNRC4, and NbPtr1. The alignment shows the first 21 amino acids of AtZAR1, NbNRC4, and NbPtr1. Blue-labeled residues from AtZAR1 and NbNRC4 indicate three hydrophobic residues from N-terminal helices that are important for triggering immunity^38,39^. Hydrophobic residues from NbPtr1 that align similarly with NbNRC4 Leu9, Leu13, and Leu19 are labeled in blue^39^. Ile13, Val21, and Val24 of NbPtr1 were mutated to glutamate, and the mutagenized residues are labeled as red in the NbPtr1 EEE mutant sequence.

(B) NbPtr1 EEE mutant and NbPtr1 K202R mutant lose autoactivity in *N. benthamiana*. EV, NbPtr1 wild-type, NbPtr1 EEE mutant, and NbPtr1 P-loop mutant (K202R) were agroinfiltrated at OD_600_ = 0.1 in *N. benthamiana* Nb-0. PCD was observed and scored at 3 dpi. Representative photographs from three independent repeats are shown in the figure. A border surrounding each photograph was colored according to the mean PCD score: black (PCD < 2) or red (PCD > 5). The scoring data of three independent repeats is represented as a bar graph with the mean PCD score + SD. GRAPHPAD PRISM v.10.1.0 was used to perform the Kruskal-Wallis test, followed by Dunn’s multiple comparisons test (*P* < 0.05), to test if the mean rank of PCD scores of NbPtr1 wild-type or mutants is significantly different from that of EV. PCD graph bars are labeled with asterisks to indicate significant differences: ****, *P* < 0.0001; ***, *P* < 0.001.

**Supplementary Figure 10. The cavity-like structure on NbPtr1 is predicted to interact with the Thr26 residue of NbNOI8.**

(A) The NbPtr1 and NbNOI-8 NOI domain interacting structure was predicted using AlphaFold3 using a seed value of 2 (Fig. 5A). This figure shows the NbPtr1 structure from the complex, with the NbPtr1-NbNOI-8 interface colored in cyan. The zoomed-in view of the cavity-like structure from NbPtr1 that holds the NbNOI-8 Thr28 residue is shown on the right.

(B) NbNOI-8 NOI domain from the interacting complex is shown in this figure (Fig. 5A). NbNOI-8 is colored cyan, and the NbPtr1-NbNOI-8 interface is colored pink. The Thr26 residue protrudes from the surface, and this structure is crucial for the interaction with NbPtr1.

**Supplementary Figure 11. Programmed cell death scoring and protein expression in Fig. 5C**

(A) Mutations on the conserved threonine residue on NbNOIs disrupt their ability to suppress NbPtr1. The scoring data of Fig. 5C are represented in a bar graph, with the mean PCD score + SD. GRAPHPAD PRISM v.10.1.0 was used to perform the Kruskal-Wallis test, followed by Dunn’s multiple comparisons test (*P* < 0.05), to test if the mean rank of PCD scores from NbNOI mutant variants is significantly different from the mean rank of PCD scores from NbNOI wild-type. PCD graph bars are labeled with asterisks to indicate significant differences: ****, *P* < 0.0001; ***, *P* < 0.001.

(B) N-terminally 1xMYC-tagged NbNOI wild-type and mutant variants were well expressed in *N. benthamiana*. EV, NbNOI-1, NbNOI-1 mutant variants (T26A, T26D, T26I), NbNOI-4, NbNOI-4 mutant variants (T28A, T28D, T28I), NbNOI-8, and NbNOI-8 mutant variants (T26A, T26D, T26I) were expressed at OD_600_ = 0.4 with P19 (OD_600_ = 0.1) in Nb-1 *ptr1*.

**Supplementary Figure 12. Mutagenesis on E431, P433, and V477 residues of NbPtr1 does not affect the autoactivity of NbPtr1.**

(A) EV, NbPtr1 wild-type and mutant variants (E431A, P433A, E431A P433A, P433E, and V477E) were expressed at OD_600_ = 0.2 in Nb-1 *ptr1*. PCD was scored and photographed at 2-3 dpi, and representative photographs from three independent repeats are shown in the figure. A border surrounding each photograph was colored according to the mean PCD score: black (PCD < 2) or red (PCD > 5).

(B) The scoring data of three independent repeats is represented as a bar graph with the mean PCD score + SD. GRAPHPAD PRISM v.10.1.0 was used to perform the Kruskal-Wallis test, followed by Dunn’s multiple comparisons test (*P* < 0.05) to test if the mean rank of PCD scores of NbPtr1 and its mutant variants is significantly different from the mean rank of PCD scores of EV. PCD bar graphs are labeled with asterisks to indicate significant differences: ****, *P* < 0.0001.

**Supplementary Figure 13. Programmed cell death scoring and ion leakage measurement of Fig. 5D**.

(A) The scoring data from three independent repeats of Fig. 5D is represented as a bar graph with the mean PCD score + SD. GRAPHPAD PRISM v.10.1.0 was used to perform the Kruskal-Wallis test, followed by Dunn’s multiple comparisons test (*P* < 0.05), to test if the mean rank of PCD scores of NbNOIs with NbPtr1 variants is significantly different from the mean rank of PCD scores of EV with NbPtr1 variants. If there is no significant difference, “ns” is indicated in the graph. PCD graph bars are labeled with asterisks to indicate significant differences: ****, *P* < 0.0001; ***, *P* < 0.001; **, *P* < 0.01; *, *P* < 0.1.

(B) Ion leakage was measured three times independently from the infiltrations described in Fig. 5D. Infiltration samples were subjected to ion leakage measurement at 3 dpi. The representative ion leakage data from three repeats are presented as a bar graph showing mean + SD. GRAPHPAD PRISM v.10.1.0 was used to perform an ordinary one-way ANOVA test, followed by Dunn’s multiple comparisons test (*P* < 0.05), to test if ion leakage from coexpression of NbNOI with NbPtr1 variants is significantly different from coexpression of EV with NbPtr1 variants. If there is no significant difference, “ns” is indicated in the graph. The graph bars are labeled with asterisks to indicate significant differences: ****, *P* < 0.0001; ***, *P* < 0.001; **, *P* < 0.01; *, *P* < 0.1.

**Supplementary Figure 14. NbNOIs and NbPtr1 variants protein expression**

EV, N-terminally 1xMYC-tagged NbNOI-1, NbNOI-4, NbNOI-6, NbNOI-8, and NbNOI-9 (OD_600_ = 0.4) were coexpressed with C-terminally 4xMYC-tagged NbPtr1 wild-type and mutant variants (E431A, P433A, E431A P433A, P433E, V477E) (OD_600_ = 0.05) in Nb-1 *ptr1*.

Due to the protein size difference of NbNOIs and NbPtr1, 8% gels were used for separating NbPtr1 proteins, and 15% gels were used for separating NbNOIs. Total protein extracts were immunoblotted with anti-MYC antibodies.

**Table S1. Sequence information of *Nicotiana benthamiana* NOI family proteins.**

**Table S2. Programmed cell death scoring data for Figure 1B. Table S3. Programmed cell death scoring data for Figure S5.**

**Table S4. Sequence information of the synthesized constructs used in this study. Table S5. Sequence information of ETI inducers used in this study.**

**Table S6. Programmed cell death scoring data for Figure 3B. Table S7. Programmed cell death scoring data for Figure S8. Table S8. Programmed cell death scoring data for Figure S9.**

**Table S9. Identifying NbPtr1 residues that interact with the conserved threonine of NbNOIs using AlphaFold3.**

**Table S10. Programmed cell death scoring data for Figure 5C, S11. Table S11. Programmed cell death scoring data for Figure S12.**

**Table S12. Programmed cell death scoring data for Figure 5D, S13A. Table S13. Oligonucleotides used in this study.**

**Table S14. Vectors used in this study.**

